## Supplementary Information for "LinguaPhylo: a probabilistic model specification language for reproducible phylogenetic analyses"

| Function | Description | Examples |
|---|---|---|
| binaryRateMatrix | Binary trait rate matrix | errorModel1.lphy, errorModel2.lphy |
| f81 | F81 model[3] | f81Coalescent.lphy |
| generalTimeReversible | General time reversible rate matrix | h5n1.lphy |
| gtr | GTR model[17] | gtrCoalescent.lphy |
| hky | HKY model[5] | hkyCoalescent.lphy |
| jukesCantor | Jukes-Cantor model[8] | jcCoalescent.lphy |
| k80 | K80 model[10] | |
| lewisMK | LewisMK model[11] | lewisMKCoalescent.lphy |
| migrationMatrix | Population process rate matrix | simpleStructuredCoalescent.lphy |
| wag | WAG model[18] | wagCoalescent.lphy |

**Table S1**: Functions for substitution models and rate matrices in LPhy.

| Generative distribution | Description | Examples |
|---|---|---|
| MultispeciesCoalescent | Multispecies coalescent | simpleMultispeciesCoalescent.lphy, simpleMultispeciesCoalescentTaxa.lphy, twoGeneMultispeciesCoalescent.lphy |
| Coalescent | Kingman's coalescent [13] | RSV2.lphy |
| SkylineCoalescent | Skyline coalescent [2] | hcv_col.lphy |
| StructuredCoalescent | Structured coalescent[12] | simpleStructuredCoalescent.lphy |

**Table S2**: Coalescent tree generative distributions in LPhy.

| Generative distribution | Description | Examples |
|---|---|---|
| BirthDeathSampling | Birth-death-sampling tree[15, 14] | birthDeathRhoSampling.lphy |
| BirthDeathSerialSampling | Birth-death serial sampling tree[16] | simpleBirthDeathSerial.lphy |
| BirthDeath | Calibrated birth-death[7] | simpleCalibratedBirthDeath.lphy, simpleExtantBirthDeath.lphy |
| FossilBirthDeathTree | Fossilized birth-death process[6] | simFossilsCompact.lphy |
| FullBirthDeath | Birth-death tree[9] | simpleFullBirthDeath.lphy |
| RhoSampleTree | Birth-death tree sampled from a larger tree | |
| SimBDReverse | Birth-death tree with extant and extinct species | simFossils.lphy |
| SimFBDAge | Birth-death tree with extant and extinct species sampled through time | simFBDAge.lphy |
| SimFossilsPoisson | Tree with fossils added to given tree at rate psi | simFossils.lphy |
| Yule | Yule tree[19] | simpleYule.lphy, yuleRelaxed.lphy |

**Table S3**: Birth-death tree generative distributions in LPhy.

| Generative distribution | Description | Examples |
|---|---|---|
| PhyloBrownian | Brownian motion process[4] | simplePhyloOU.lphy |
| PhyloCTMC | Continuous time Markov process[3] | simpleBModelTest.lphy |
| PhyloMultivariateBrownian | Multivariate Brownian motion | simplePhyloMultivariateBrownian.lphy |
| PhyloOU | Ornstein-Ulhenbeck process[4] | simplePhyloBrownian.lphy |

**Table S4**: Phylogenetic likelihood distributions in LPhy.

| Generative distribution | Description | Examples |
|---|---|---|
| Bernoulli | Coin toss distribution | simpleRandomLocalClock.lphy, simpleBModelTest.lphy |
| Beta | Beta distribution | birthDeathRhoSampling.lphy, simpleBModelTest.lphy |
| Cauchy | Cauchy distribution | |
| Dirichlet | Dirichlet distribution | birthDeathRhoSampling.lphy, dirichlet.lphy |
| DiscreteUniform | Discrete-uniform distribution | simpleBModelTest.lphy, simpleBModelTest2.lphy |
| DiscretizeGamma | Discretize-gamma distribution | gtrGammaCoalescent.lphy, simpleBModelTest.lphy |
| Exp | Exponential distribution | birthDeathRhoSampling.lphy, yuleRelaxed.lphy |
| ExpMarkovChain | Smoothing distribution [2] | skylineCoalescent.lphy |
| Gamma | Gamma distribution | covidDPG.lphy |
| Geometric | Geometric distribution | |
| InverseGamma | Inverse-gamma distribution | totalEvidence.lphy |
| LogNormal | Log-normal distribution | hkyCoalescent.lphy, errorModel1.lphy |
| Normal | Normal distribution | simplePhyloBrownian.lphy, simplePhyloOU.lphy |
| NormalGamma | Normal-gamma distribution | simplePhyloBrownian.lphy, simplePhyloOU.lphy |
| Poisson | Poisson distribution | expression4.lphy, simpleRandomLocalClock2.lphy |
| RandomBooleanArray | Samples a random boolean array | simpleRandomLocalClock2.lphy |
| RandomComposition | Samples a random k-tuple of positive integers that sum to n | skylineCoalescent.lphy |
| Uniform | Uniform distribution | simFossilsCompact.lphy |
| Weibull | Weibull distribution | |
| WeightedDirichlet | Weighted dirichlet distribution | totalEvidence.lphy, weightedDirichlet.lphy |

**Table S5**: Parametric distributions in LPhy.

| Function | Description | Examples |
|---|---|---|
| aminoAcids | Amino acid data type | wagCoalescent.lphy |
| binaryDataType | Binary data type | |
| nucleotides | Nucleotide data type | primates2.lphy |
| standard | Standard data type | totalEvidence.lphy |

**Table S6**: Alignment data types in LPhy.

| Function | Description | Examples |
|---|---|---|
| nucleotideModel | bModelTest[1] rate matrix | simpleBModelTest.lphy, simpleBModelTest2.lphy |
| bModelSet | bModelTest model set | simpleBModelTest.lphy |
| bSiteRates | Site rates for the given bModelTest parameters | simpleBModelTest2.lphy |
| bSiteModel | bModelTest site model | simpleBModelTest.lphy |

**Table S7**: Bayesian phylogenetic site model averaging in LPhy.

[17] S Tarvaré. Some probabilistic and statistical problems in the analysis of dna sequences. *Some mathematical question in biology-DNA sequence analysis*, 1986.

[18] Simon Whelan and Nick Goldman. A general empirical model of protein evolution derived from multiple protein families using a maximum-likelihood approach. *Molecular biology and evolution*, 18(5):691–699, 2001.

[19] George Udny Yule. Ii.—a mathematical theory of evolution, based on the conclusions of dr. jc willis, fr s. *Philosophical transactions of the Royal Society of London. Series B, containing papers of a biological character*, 213(402-410):21–87, 1925.
